# A novel nuclear speckle factor, USP42, promotes homologous recombination repair by resolving DNA double-strand break induced R-loop

**DOI:** 10.1101/776823

**Authors:** Misaki Matsui, Ryo Sakasai, Masako Abe, Yusuke Kimura, Shoki Kajita, Wakana Torii, Yoko Katsuki, Masamichi Ishiai, Kuniyoshi Iwabuchi, Minoru Takata, Ryotaro Nishi

## Abstract

The nucleus of mammalian cells is highly compartmentalized by nuclear bodies, including nuclear speckles. While nuclear bodies are known to function in regulating gene expression, their involvement in DNA repair has not been actively investigated. Here, our focused screen for nuclear speckle factors involved in homologous recombination (HR), which is a faithful DNA double-strand break (DSB) repair mechanism, revealed that nuclear speckle factors regulating transcription are potentially involved in the regulation of HR. Among the top hits, we provide evidence showing that USP42, which is a deubiquitylating enzyme and a hitherto unidentified nuclear speckles factor, promotes HR by facilitating BRCA1 recruitment to DSB sites and DNA-end resection. We further showed that USP42 localizes to nuclear speckles via an intrinsically disordered region, which is required for efficient HR. Furthermore, we established that USP42 interacts with DHX9, which possesses DNA-RNA helicase activity, and is required for efficient resolution of DSB-induced R-loop. Mechanistically, USP42 antagonizes mono-ubiquitylation of DHX9 that is evoked after DSB induction. In conclusion, our data propose a model in which a novel nuclear speckle factor, USP42, facilitates DSB-induced R-loop resolution, BRCA1 loading to DSB sites and preferential DSB repair by HR, indicating the importance of spatial regulation of DSB repair choice mediated by nuclear bodies.

**Significant statement:** Defects in the repair of DNA double-strand break (DSB), which is one of the most harmful DNA insults, cause human diseases including cancers. It has been suggested that DSBs generated in the coding region tend to be repaired by homologous recombination (HR) that is error-free DSB repair pathway. To reveal the spatial regulation of HR, in this study, we investigated the potential contribution of nuclear bodies, especially nuclear speckles, to HR, identifying a deubiquitylating enzyme USP42 as a HR promoting factor. We found that USP42 deubiquitylates DHX9, facilitates resolution of DNA-RNA hybrid structure and enhances HR through BRCA1 loading to DSB sites.

**Classification:** Biological Sciences, Cell Biology

## Introduction

Genomic DNA of mammalian cells does not distribute uniformly in the nucleus. The nucleus of mammalian cells contains various nuclear bodies, such as promyelocytic leukaemia bodies, Cajal bodies, speckles, paraspeckles and nuclear speckles, which are membraneless structures that highly compartmentalize the nucleus (1, 2). Although these nuclear bodies play important roles in expressing genomic function, including stress responses, messenger RNA (mRNA) splicing and transcription (3), the involvement of these nuclear bodies in the regulation of DNA repair remains elusive. Nuclear speckles (also designated interchromatin granule clusters) are self-organizing membraneless nuclear bodies that are detected as 20 to 50 irregularly shaped dots and localized in spaces between chromatins (4). *In situ* hybridization and immunofluorescence staining of a representative nuclear speckle factor, SC35, revealed that nuclear speckles frequently localized next to transcriptionally active gene loci (5-9). In accordance with their subnuclear localization, nuclear speckles are suggested to function as storage sites of mRNA splicing factors and transcription factors (10-12). In addition, some groups have suggested a direct contribution of nuclear speckles to splicing and transcriptional regulation (11, 13-15). Furthermore, in addition to poly(A)^+^ RNAs and non-coding RNAs (16, 17), mass spectrometry analysis of purified mouse liver nuclear speckles indicated that nuclear speckles are composed of not only proteins involved in RNA metabolism but also factors contributing to other cellular functions, such as apoptosis (18, 19), suggesting that nuclear speckles may play roles in a wide variety of cellular functions.

DNA double-strand breaks (DSBs) are likely the most cytotoxic DNA insults generated by endogenous sources and exogenous reagents such as ionizing radiation (IR). It is well known that dysfunctions in the DSB repair machinery cause human hereditary diseases, which often feature predisposition to tumorigenesis, immune deficiencies and cellular hypersensitivity to IR, highlighting the importance of DSB repair for maintaining individual wellness and genome integrity (20, 21). DSBs are repaired mainly by two mutually exclusive pathways: non-homologous end-joining (NHEJ) and homologous recombination (HR). In contrast to error-permissive NHEJ, HR is thought to be an error-free pathway since HR copies the DNA sequence from the undamaged sister chromatid in most cases (22, 23). It has been reported that DSBs generated in actively transcribed genes tend to be repaired by HR (24, 25), suggesting that HR could be preferentially chosen to accurately preserve genetic information for coding regions. In this study, we investigated the potential regulatory mechanism of HR mediated by nuclear speckles, which are also functionally associated with RNA metabolism.

## Results

### Nuclear speckle factor screening for HR regulation indicated a cross-talk between HR and transcription

To investigate whether nuclear speckles spatially contribute to proper DSB responses, especially HR, camptothecin (CPT)-induced phosphorylation of RPA2 on Ser4 and Ser8 (pRPA2 S4/S8), which is thought to be an indicative of the early HR process, was examined in the presence of tubercidin, which induces the dispersion of some nuclear speckle factors, including SRSF1, SC35 and poly(A)^+^ RNA (Fig. 1*A*) (26). As shown in Fig. 1*B*, tubercidin treatment resulted in reduced phosphorylation of RPA2, suggesting that the integrity of nuclear speckles or the localization of some nuclear speckle factors to distinct foci is required for efficient HR. To evaluate the effect of nuclear speckle factors on HR, we investigated the frequency of non-crossover gene conversion-mediated HR with a Direct-Repeat GFP (DR-GFP) assay by knocking down nuclear speckle factors. A short interfering RNA (siRNA) library comprised 129 siRNAs targeting potential nuclear speckle factors identified by proteomics analysis, while proteins involved in mRNA splicing were excluded because of a preciously suggested intervention for DSB repair (27). The siRNA library also included siRNAs targeting the deubiquitylating enzyme USP42, which showed a dot-like localization similar to nuclear speckles (28, 29) and was found actually to colocalize with SC35 and to disperse in the nucleoplasm to a lesser extent (Fig. S1*A*). For each target, two siRNAs were pooled for transfection. The efficiencies of HR were Z-score normalized, identifying several factors that potentially regulated HR together with a positive control (siCtIP) and factors already indicated to promote HR, such as XAB2 and ZMYND8 (Fig. 1*C*) (30, 31). Furthermore, the DR-GFP assay was carried out with two individual siRNAs targeting WDR5, DHX9, GTF3C2 and USP42, verifying the results of the initial screen and identifying these factors as novel HR regulatory proteins (Fig. 1*D* and S1*B*). Since USP42 is a hitherto unidentified nuclear speckle factor and is a unique deubiquitylating enzyme that localizes to nuclear speckles, we decided to focus on USP42 to reveal a nuclear speckle-mediated HR regulation.

**Fig. 1.**
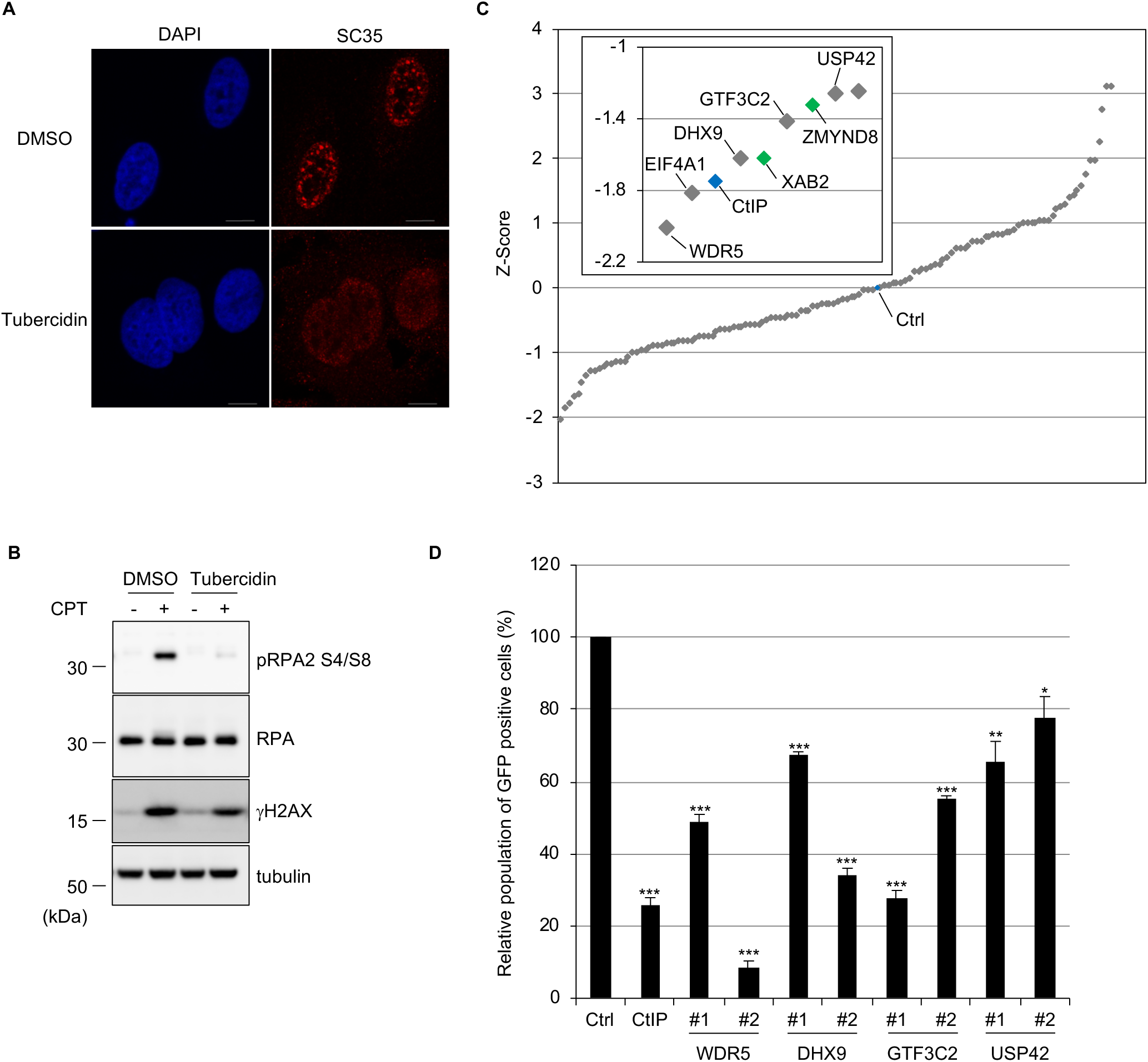
Nuclear speckle factor screening for HR regulation indicated a cross-talk between HR and transcription. (A) U2OS cells treated with 2.5 μg/ml tubercidin or mock treated (dimethyl sulfoxide: DMSO) were subjected to immunofluorescence staining with the anti-SC35 antibody. Scale bar: 10 μm. (B) U2OS cells were incubated with 2.5 μg/ml tubercidin or DMSO followed by treatment with CPT (1 μM, 1 hour). Cell extracts were subjected to immunoblotting analysis with the indicated antibodies. (C) Screen for nuclear speckle factors involved in HR regulation. The DR-GFP assay was performed with siRNA pools targeting nuclear speckle factors. The siRNA targeting CtIP or luciferase (control: Ctrl) was used as a positive or negative control, respectively, and indicated with blue markers. Homology directed repair efficiency is plotted as a Z-score. The inset magnifies the data for top hits with gene symbols. The genes previously reported to be involved in HR regulation are indicated with green. (D) A DR-GFP assay was carried out with two independent siRNAs targeting selected top hits from the initial screen. GFP-positive cell populations were normalized to mock treatment, which was set to 100% (mean ± SEM, n=3). *: p<0.05. **: p<0.01. ***: p< 0.005. Also see Fig. S1.

### USP42 promotes HR by recruiting BRCA1 to DSB sites

To further understand how USP42 promotes HR, the effects of USP42 depletion with siRNAs on various aspects of HR were investigated. Depletion of USP42 with two siRNAs resulted in reduced phosphorylation of RPA2 upon CPT treatment, suggesting that USP42 could function in early phases of HR (Fig. 2*A*). Thus, we examined DNA-end resection that generates 3’ single-stranded DNA overhangs by nucleolytic degradation of the 5’ terminated stand of the DSB upon CPT treatment, indicating that USP42 is required for efficient DNA-end resection (Fig. 2*B*). Since DNA-end resection also occurs during single-strand annealing and break-induced replication, which are RAD51-independent DSB repair pathways, RAD51 foci formation was tested. Depleting USP42 resulted in significantly reduced RAD51 foci formation, suggesting that USP42 functions upstream of RAD51 and promotes HR rather than RAD51-independent DSB repair pathways (Fig. 2*C* and S2*A*). These results are in good agreement with DR-GFP assay-based screen data, which were not due to smaller population in S and G2 phase (Fig. 1*D*, S2*B* and S2*C*). Furthermore, USP42 depletion sensitized cells to IR, indicating its physiological importance (Fig. 2*D*). In addition, USP42 depletion did not affect the protein levels of HR factors involved in DNA-end resection, such as the Mre11-Rad50-Nbs1 complex and CtIP (Fig. S2*D*). Furthermore, to carry out rescue experiments, we created a USP42 knockout U2OS cell line (USP42 KO). This cell line recapitulated the phenotypes caused by USP42 knockdown (Fig. S2*E* and S2*F*), which were rescued by stable expression of exogenous GFP-tagged USP42 but not of GFP (Fig. 2*E* and 2*F*). As DSBs generated in transcriptionally active regions tend to be repaired by HR, we speculated that USP42, which localizes close to actively transcribed loci, may favour HR over NHEJ for these DSBs. Supporting this idea, knockdown of USP42 resulted in reduced BRCA1 foci, which promotes HR, in CENPF-positive cells, whereas the number of 53BP1 foci per cell, which counteracts BRCA1 loading, was increased (Fig. 2*G*, 2*H* and S2*G*). Altogether, these results suggest that USP42 facilitates BRCA1 recruitment to DSB sites and consequently promotes DNA-end resection, RAD51 loading and HR.

**Fig. 2.**
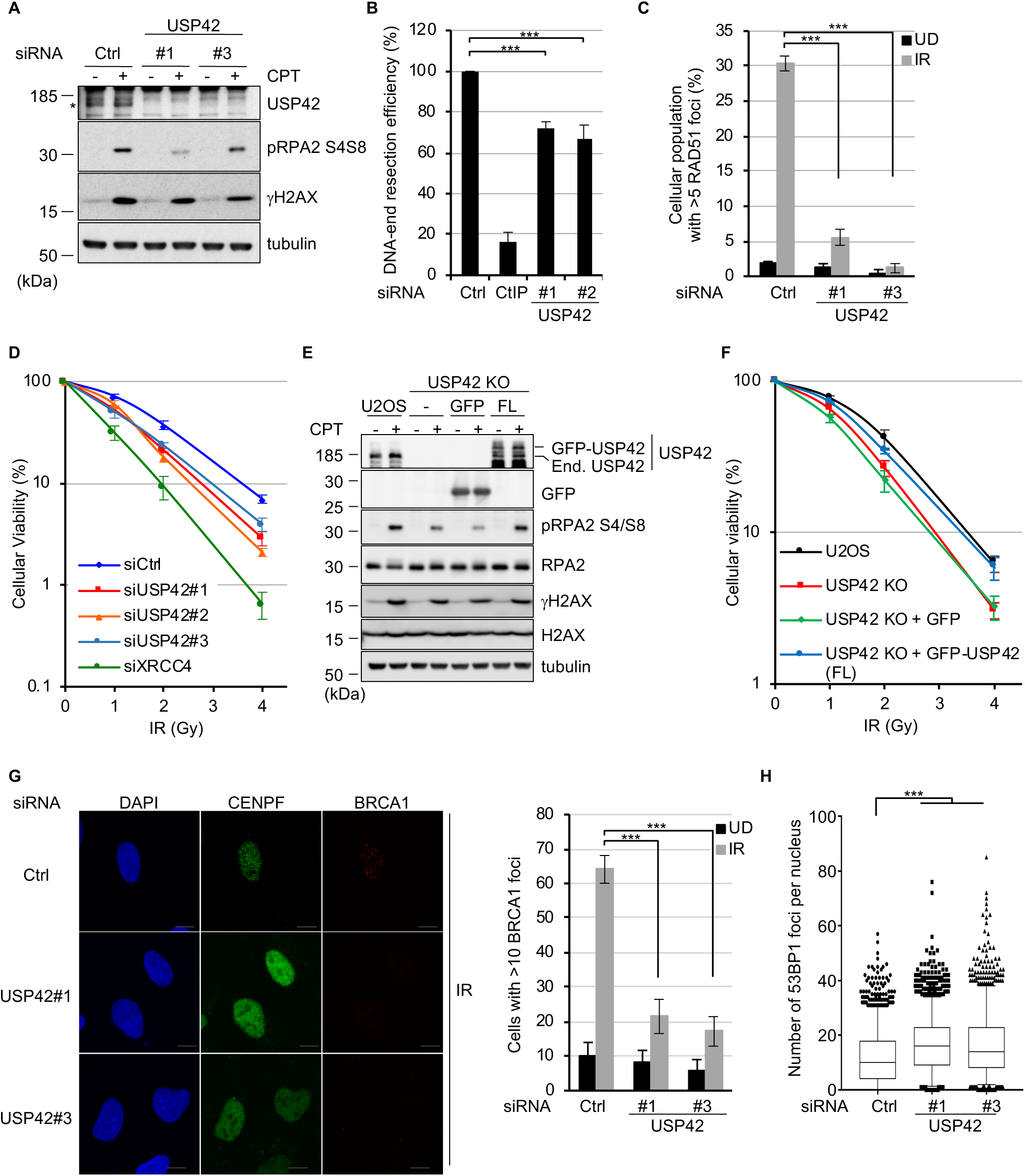
USP42 promotes HR by recruiting BRCA1 to DSB sites. (A) U2OS cells transfected either with siRNA control (Ctrl) or siRNA targeting USP42 were treated with CPT or mock treated. Cell extracts were analysed by immunoblotting with the indicated antibodies. Asterisk indicates endogenous USP42. Immunoblotting with the anti-γH2AX antibody was used as a control for DNA damage induction. (B) A DNA-end resection assay was carried out with the cells transfected with the indicated siRNAs (mean ± SEM, n=3). The siRNA targeting CtIP was a positive control. (C) RAD51 foci formation efficiency was examined with cells transfected with the indicated siRNAs (mean ± SEM, n=3). The population of cells with >5 RAD51 foci was plotted. (D) Cells transfected with the indicated siRNAs were subjected to a clonogenic survival assay (mean ± SEM, n≥3). The siRNA targeting XRCC4 was used as a positive control. (E) U2OS, USP42 KO or USP42 KO cells complemented with either GFP or GFP-USP42 (FL) were treated with CPT and subjected to immunoblotting analysis with the indicated antibodies. End. USP42: endogenous USP42. (F) The indicated cell lines were subjected to a clonogenic survival assay (mean ± SEM, n≥4). (G) BRCA1 foci formation efficiency was examined with cells transfected with the indicated siRNAs. (Left) The representative images are shown. (Right) The population of cells with >10 BRCA1 foci was plotted (mean ± SEM, n=3). The cells positive for CENPF staining were analysed. See Fig. S2*G* for the representative images of undamaged condition. (H) 53BP1 foci formation efficiency was examined with cells transfected with the indicated siRNAs. The number of foci per nucleus after IR were plotted as a box and whiskers plot (median, 5 to 95 percentile, n=3). The cells positive for CENPF staining were analysed. ***: p< 0.005. See also Fig. S2.

### Nuclear speckle localization of USP42 is required for efficient HR

In ensuing studies, we examined whether the subnuclear localization of USP42 plays a role in its function in HR by elucidating the nature of the nuclear speckle localization of USP42. As shown in Fig. 3*A*, except for the ubiquitin-specific protease (USP) domain that is a deubiquitylating domain, USP42 does not contain obvious functional domains, whereas proline (P)-, arginine (R)- and lysine (K)-rich regions are found in its carboxyl-terminal half. First, we asked whether the enzymatic activity of USP42 is required for its nuclear speckle localization (Fig. 3*A* and 3*B*). Whle the USP42 mutant lacking the USP domain (ΔUSP) colocalized with SC35, the ΔC mutant (1-412 amino acids (a. a.) residues) that included the USP domain showed dispersed nuclear and partial cytoplasmic localization, suggesting that the enzymatic activity of USP42 is dispensable for its nuclear speckle localization. To identify the domain(s) responsible for nuclear speckle localization, the localization of various truncated mutants of USP42 was examined (Fig.3*A* and 3*B*). USP42 lacking the C-terminal region (1-945 a. a.) almost completely localized to the cytoplasm, whereas the C-terminal region (946-1316 a. a.) was sufficient for nuclear and nuclear speckle localization. Deletion mutants of this C-terminal region revealed that 211 (946-1156 a. a.) and 160 residues (1157-1316 a. a.) functioned as a nuclear speckle localization signal domain, although the former region represented clearer colocalization with SC35. Further segmentation of the former domain resulted in a diffuse and large dot-like pattern that was not likely to be nuclear speckles (Fig. S3*A* and S3*B*), suggesting that the integrity of this region is important for nuclear speckle localization of USP42. Deleting these regions from the USP42 protein revealed that a region from 946 to 1196 residues is required for nuclear speckle localization of USP42. This region weakly matched the arginine/serine (RS) repeat motif, which was previously reported as a nuclear speckle localization signal (Fig. S3*C*) (32-34). Since it is suggested that the intrinsically disordered region (also known as the low complexity domain) plays a pivotal role in the formation of membraneless organelles by liquid-liquid phase separation (LLPS) (35), USP42 was analysed for an intrinsically disordered region (Fig. S3*D*) (36). As expected, USP42 was predicted to be mostly disordered, including a nuclear speckle localization signal (946-1196 a. a.). Finally, we examined CPT-induced signalling in USP42 KO cell lines that stably express either GFP or GFP-fused USP42 (full-length: FL, ΔC or Δ946-1196 a. a.) (Fig. 3*C* and 3*D*). Although exogenously expressed GFP-USP42 (FL) restored CPT-induced phosphorylation of RPA2, cells expressing truncated mutants of USP42 that lost nuclear speckle localization failed to rescue this phenotype, suggesting that nuclear speckle localization of USP42 is indispensable for proper HR progression.

**Fig. 3.**
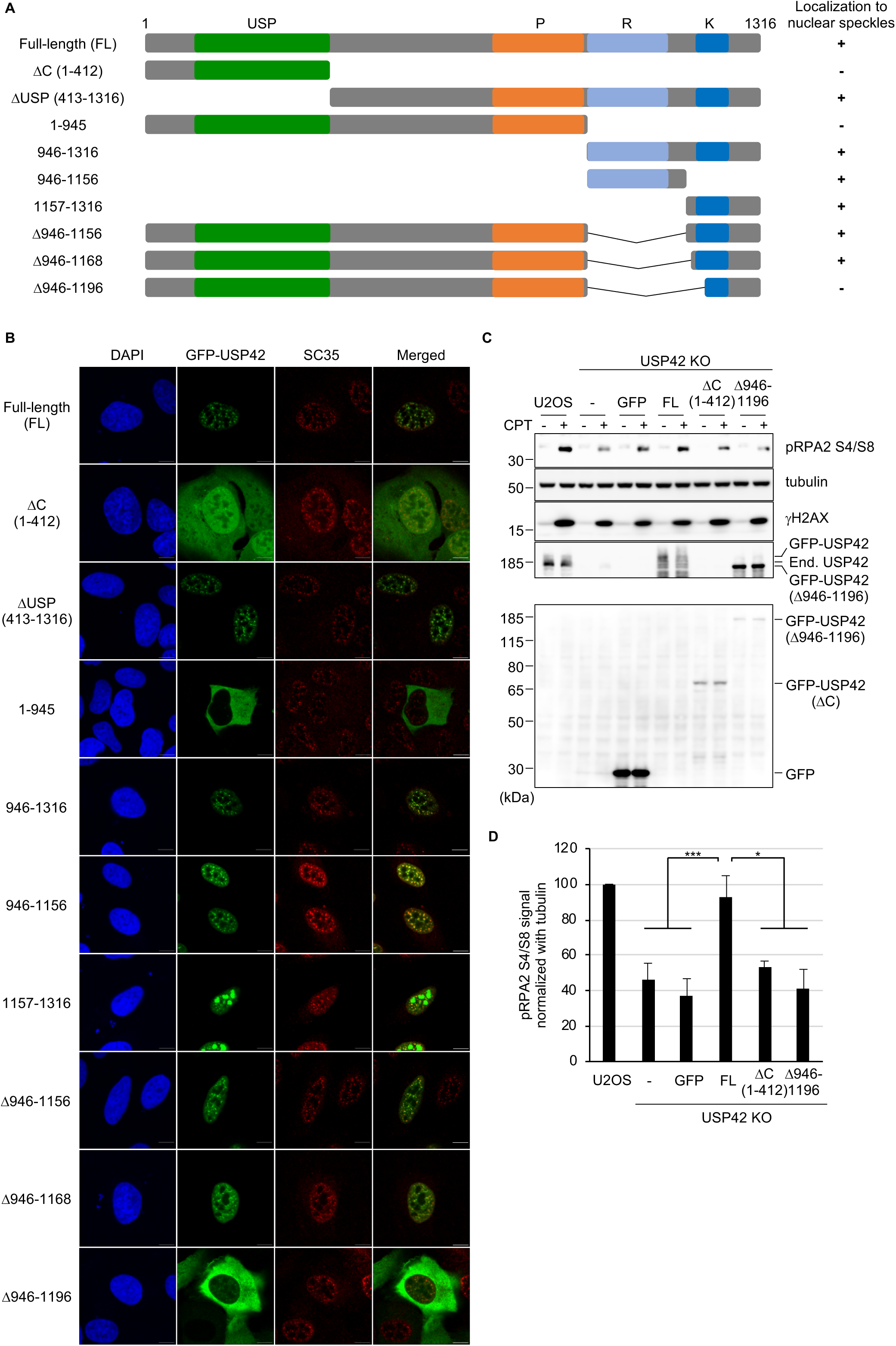
Nuclear speckle localization of USP42 is required for efficient HR. (A) Schematic representation of USP42 protein and truncated mutants. The numbers represent amino acid residues. USP: ubiquitin-specific protease domain. P: proline-rich region. R: arginine-rich region. K: lysine-rich region. Localization to nuclear speckles of the indicated mutants are shown on the right. (B) Subcellular localization of various GFP-fused USP42 proteins. Cells transfected with the plasmids encoding the indicated GFP-USP42 were subjected to immunofluorescence staining with an anti-SC35 antibody. Scale bar: 10 μm. (C) U2OS or USP42 KO cells complemented with either GFP or GFP-USP42 (FL, ΔC or Δ946-1196) were treated with CPT (1 μM, 1 hour) and subjected to immunoblotting analysis with the indicated antibodies. Note that endogenous USP42 (End. USP42) and GFP-USP42 (FL) could only be detected in pellet fraction with an anti-USP42 antibody, while GFP and GFP-USP42 (ΔC) were detected in soluble fraction with an anti-GFP antibody. GFP-USP42 (Δ946-1196) was mainly detected in the pellet fraction. (D) The signal intensities of phosphorylated RPA2 that were normalized to the signal intensity of tubulin were plotted (mean ± SEM, n≥3). *: p<0.05. ***: p< 0.005. See also Fig. S3.

### USP42 is epistatic with DHX9 in the cellular survival after DSB induction and promotes resolution of DSB-induced R-loop

Since USP42 is a deubiquitylating enzyme, we assumed that USP42 may fulfil its function by deubiquitylating interacting partner(s). Thus, mass spectrometry analysis of affinity-purified GFP-USP42 interacting proteins was performed with a negative control (i.e., GFP), identifying 161 proteins that specifically interact with USP42 (data not shown). Focusing on DHX9 (also known as RHA or NDHII), which is one of the top hits and belongs to the DEAH RNA helicase family, we confirmed that USP42 interacted with DHX9 in a reciprocal manner by coimmunoprecipitation (Fig. 4*A* and 4*B*). To investigate whether DHX9 is involved in HR, we assessed CPT-induced phosphorylation of RPA2 with two independent siRNAs targeting DHX9. Depletion of DHX9 resulted in a reduced pRPA2 S4/S8 signal (Fig. 4*C*), which was similar to USP42 depletion (Fig. 2*E* and S2*E*). In addition, depleting DHX9 resulted in impaired HR activity with the DR-GFP assay (Fig. 1*D* and S4*A*) without affecting the cell cycle profile (Fig. S4*B*). These findings prompted us to examine whether USP42 and DHX9 function in the same axis in HR. While depletion of DHX9 in U2OS cells resulted in increased sensitivity to IR compared to control siRNA, knocking down DHX9 did not confer further sensitivity to IR in USP42 KO cells (Fig. 4*D*), indicating that USP42 is epistatic with DHX9, and both proteins are required for proper HR. Since it was previously suggested that DHX9 promotes DNA-RNA hybrid structure (R-loop) resolution *in vivo* and *in vitro* (37-39), we speculated that USP42 could facilitate R-loop resolution together with DHX9 in HR. In line with this idea, the signal intensity of the S9.6 antibody that specifically detects the R-loop was increased upon DSB induction and remained increased in USP42 KO cells compared to U2OS cells (Fig. 4*E*), indicating that USP42 is required for DSB-induced R-loop resolution. Contrary, without DSB-induction, the signal intensity of the S9.6 antibody was significantly decreased in USP42 KO cells compared to U2OS cells (Fig. S4*C*), suggesting DSB-dependent and DSB-independent R-loop regulation.

**Fig. 4.**
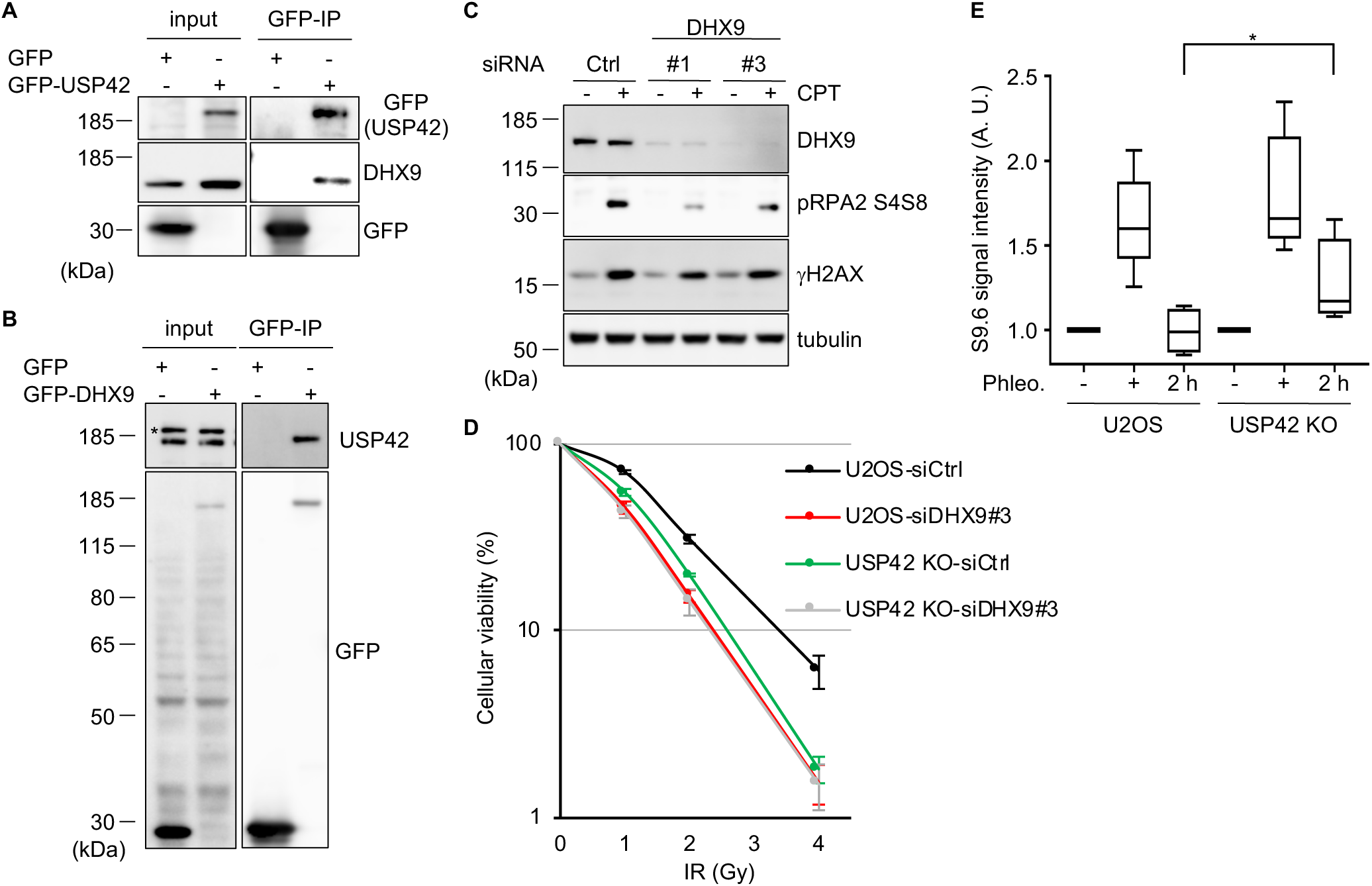
USP42 is epistatic with DHX9 in the cellular survival after DSB induction and promotes resolution of DSB-induced R-loop. (A, B) The interaction between USP42 and DHX9 was tested by coimmunoprecipitation with either GFP-USP42 (A) or GFP-DHX9 (B) followed by immunoblotting analysis with the indicated antibodies. Transfection with a plasmid encoding GFP was a negative control. (C) U2OS cells transfected with the indicated siRNAs were treated with CPT and then subjected to immunoblotting analysis with the indicated antibodies. (D) U2OS or USP42 KO cells transfected with the indicated siRNAs were subjected to a clonogenic survival assay (mean ± SEM, n≥4). (E) The indicated cells were treated with phleomycin (Phleo., +) and then further cultured for 2 hours upon removal of phleomycin (2 h). Purified genomic DNA was subjected to slot blot analysis with an S9.6 antibody. Signal intensity was normalized to mock treated samples of each cell lines (-), and then plotted as a box and whiskers plot (median, 5 to 95 percentile, n=4). *: p<0.05. See also Fig. S4.

### USP42 antagonizes mono-ubiquitylation of DHX9 that is evoked after DSB induction

To further examine how USP42 contribute to R-loop resolution and HR, the effect of USP42 depletion on the potential ubiquitylation of DHX9 was tested. Ubiquitylation of transiently over-expressed GFP-tagged DHX9 (GFP-DHX9) was detected in parental U2OS cells, which was pronounced in USP42 KO cells (Fig. 5*A*). In line with this notion, over-expressed HA-tagged ubiquitin preferentially immunoprecipitated endogenous DHX9 in USP42 KO cells in comparison to U2OS cells (Fig. 5*B*). Furthermore, it is worth noting that ubiquitylation of DHX9 was enhanced upon CPT treatment in U2OS cells, while in USP42 KO cells, ubiquitylation of DHX9 was maintained at increased levels regardless of CPT treatment (Fig. 5*A* and *B*), suggesting that mono-ubiquitylation of DHX9, which is suppressed by USP42 when cells are not with DSB, is induced only after DSB induction. Thus, we propose a model in which USP42 and DHX9 function to resolve the R-loop generated near DSBs and nuclear speckles, which facilitates BRCA1 loading to sites of damage and preferential DSB repair by HR (Fig. 5*C*).

**Fig. 5.**
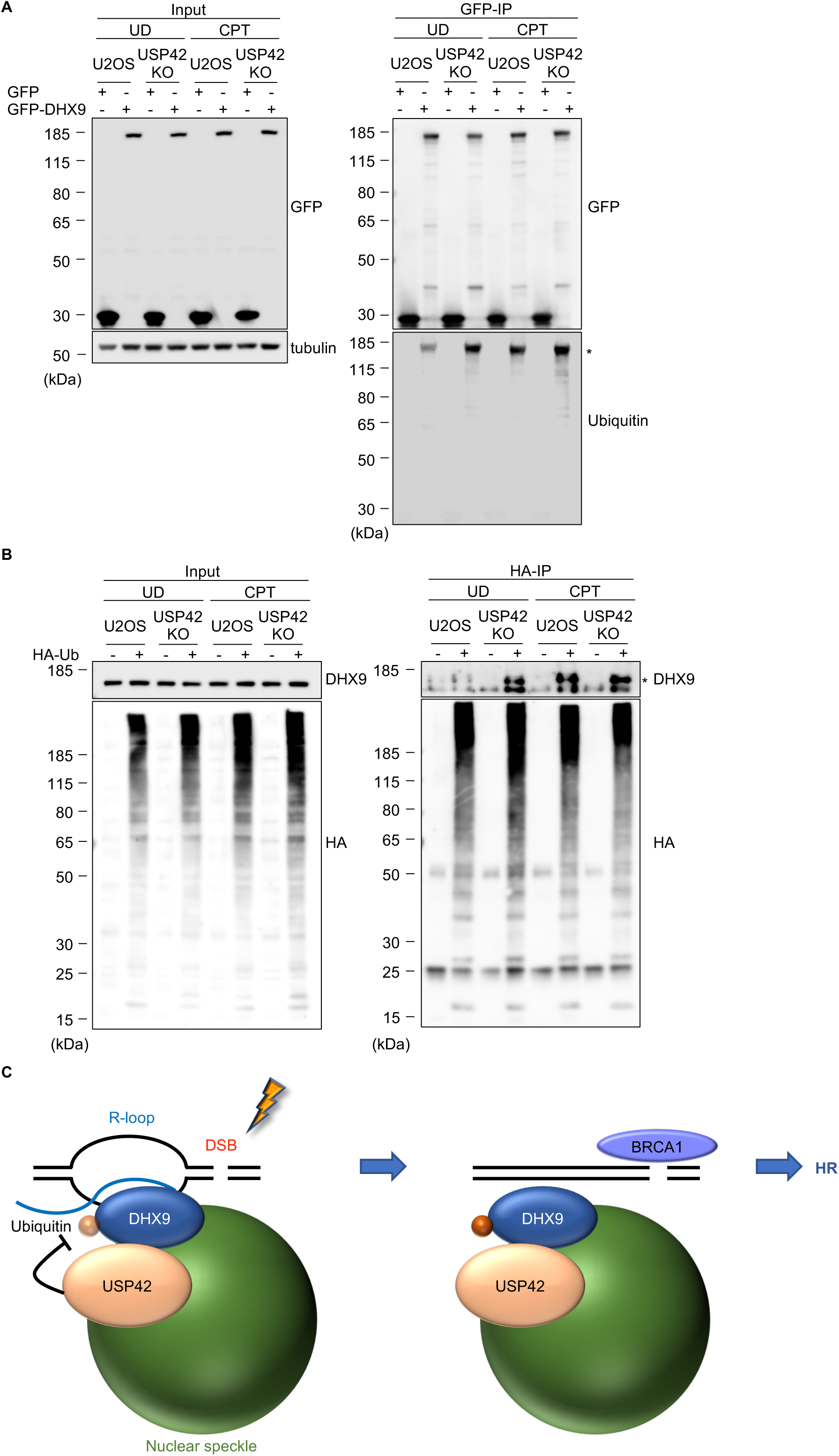
USP42 antagonizes mono-ubiquitylation of DHX9 that is evoked after DSB induction. (A, B) U2OS or USP42 KO cells transfected with GFP-DHX9 (A) or HA-tagged ubiquitin (B) were treated with CPT or mock treated. Immunoprecipitation with either anti-GFP (A) or anti-HA (B) antibodies was followed by immunoblotting with the indicated antibodies. Asterisks indicate monoubiquitylated DHX9. (C) A model proposed by our study. See main text for the details.

## Discussion

Our focused screen with a DR-GFP reporter assay identified several nuclear speckle factors potentially involved in HR regulation. Among these were transcription regulators, including WDR5, ZMYND8 and USP42. It is well known that WDR5 forms the histone H3 lysine 4 methyltransferase complex with mixed lineage leukaemia (MLL, MLL1-4) and promotes RNA polymerase II-dependent transcription activation (40). In addition, it was recently suggested that WDR5, BRCA1 and BRAD1 were functionally connected to suppress DNA damage during reprogramming (41). ZMYND8 was reported to play roles in activating transcription and promoting HR via multivalent binding to chromatin, including histone H4 acetlylations (30, 42, 43). In this study, we revealed that USP42, which was previously suggested to enhance transcription by targeting histone H2B ubiquitylation (28), is required for efficient HR. These findings validating our screen results might suggest that active transcription initiation or elongation in the vicinity of nuclear speckles is required for efficient HR or favouring HR over NHEJ, consistent with the idea termed transcription-associated HR (44). To further understand the spaciotemporal regulation of HR, it will be essential to reveal the underlying molecular mechanisms of these proteins, which are associated with different chromatin modifications. It is worth noting that GTF3C2, which is a subunit of transcription factor TFIIIC and plays roles in the expression of non-coding RNAs such as human Alu RNA and mouse B2 RNA by RNA polymerase III, was identified in our screen. These non-coding RNAs were upregulated by stresses, including etoposide and IR (45), and inhibited transcription by RNA polymerase II (46, 47). Therefore, it will be an exciting research area to explore non-coding RNA-mediated transcription inhibition and DSB repair regulation mediated by nuclear speckles.

Through domain mapping analysis of USP42, an intrinsically disordered region (946 to 1196 a. a.) was identified as a signal domain for nuclear speckle localization, which is important for HR promotion. Interestingly, DHX9, identified as a USP42 interactor, is also predicted to contain another type of disordered domain, the prion-like domain (1169-1270 a. a.) that is suggested to play roles in phase separation (48). Moreover, a USP42 mutant that interacted with DHX9 but lost nuclear speckle localization failed to rescue defective HR caused by USP42 KO (data not shown), suggesting that locally increased concentrations of both proteins are important for proper HR. LLPS that is driven by intrinsically disordered domains was previously implicated in transcriptional regulation by increasing local concentrations of proteins and compartmentalization (35); thus, our data may suggest another biological function, DSB repair, which is regulated by LLPS. It had also been suggested that LLPS was regulated by RNA binding to intrinsically disordered regions such as RS repeats (49). Therefore, it will be interesting to examine whether LLPS mediated by RNA, including non-coding RNAs, contributes to DNA repair.

In the present research, we indicate that USP42 promotes HR by facilitating resolution of the DSB-induced R-loop, recruiting BRCA1 to DSB sites and promoting DNA-end resection. Furthermore, ubiquitylation of DHX9 was antagonized by USP42 in the absence of DSB (Fig. 5*A* and *B*), suggesting that DHX9 ubiquitylation may be important for the activation of DHX9. Although, consistent with this hypothesis, R-loop was decreased in USP42 KO cells without DSB induction in which DHX9 is constantly ubiquitylated (Fig. S4*C*), how DHX9 distinguishes DSB-induced R-loop an “normal” R-loop should be addressed in the future works. On the other hand, depleting USP42 or its interacting partner DHX9 resulted in mildly increased sensitivity to IR, suggesting that only a subset of DSB repair, likely HR, required these proteins. Although it is evident that processing of the R-loop could positively and negatively influence DSB repair (50), how the R-loop affects DSB repair might be dependent on the stage of DSB repair, transcriptional activity and the genomic location of DSB. Such complexity of R-loop-mediated regulation of DSB repair can explain why various proteins, including RAD52-XPG and SETX, have been implicated in R-loop resolution (44, 51, 52).

## Materials and methods

### Cell lines and cell culture

All cell lines were cultured at 37°C in a humidified 5% CO_2_ atmosphere. U2OS cells were cultured with Dulbecco’s modified Eagle’s medium (DMEM, Nacalai tesque) containing 10% fetal bovine serum (FBS, SIGMA), 100 U/ml penicillin (Nacalai tesque), 100 μg/ml streptomycin (Nacalai tesque) and 584 μg/ml L-glutamine. The U2OS USP42 KO cell line and its complemented cell lines stably expressing either GFP or GFP-USP42 (FL, ΔC or Δ946-1196) were cultured with identical media with 1 μg/ml puromycin (InvivoGen) or with 1 μg/ml puromycin and 500 μg/ml Geneticin, respectively.

### Transfection of plasmids and siRNAs

Cells were transfected with the plasmids or siRNAs (40 nM at final concentration) by Mirus TransIT-LT1 (Mirus Bio LLC) or HiperFect (Qiagen) according to the manufacturer’s instructions, respectively. See Table S1 for the oligonucleotides used in this study.

### DR-GFP assay

The DR-GFP assay was carried out as previously described (29). HR repair efficiency was investigated by a chromosomal DSB-induced gene conversion assay system with transient expression of the I-SceI restriction enzyme in U2OS cells carrying a Direct-Repeat GFP reporter as previously described (53).

### Cell extract preparation and immunoblotting analysis

Except for sample preparation for mass spectrometry analysis and immunoprecipitation, cell extracts were prepared with CSK buffer [10 mM PIPES (pH 6.8), 3 mM MgCl_2_, 1 mM ethylene glycol tetraacetic acid (EGTA), 0.1% Triton X-100 and 300 mM sucrose] containing 300 mM NaCl, 1 x Protease Inhibitor cocktail ethylenediaminetetraacetic acid (EDTA)-free (PI, Roche), 10 mM NaF (Nacalai tesque), 20 mM N-ethylmaleimide (NEM, Nacalai tesque) and 0.25 mM phenylmethylsulfonyl fluoride (PMSF, Sigma-Aldrich). The cells were washed twice with ice-cold PBS and incubated with an appropriate volume of CSK buffer for 1 hour on ice with occasional mixing. Soluble fractions were collected by centrifugation at 20,000 x g for 10 min at 4°C. Residual chromatin fractions (pellet fractions) were washed twice with identical buffer and then solubilized by sonication (TOMY, UD-100, 40% output, 30 sec). Where indicated, cells were incubated with 2.5 μg/ml tubercidin (Sigma-Aldrich) for 2 hours and/or 1 μM CPT for 1 hour. For mass spectrometry analysis and immunoprecipitation, cells were washed twice with ice-cold PBS and collected with an appropriate volume of ice-cold PBS, followed by centrifugation at 10,000 x g, for 1 min. Cells were lysed with IP lysis buffer [20 mM Tris-HCl (pH 7.5), 2 mM MgCl_2_, 0.5% NP-40 and 10% glycerol] containing 40 mM NaCl, 1 x PI, 10 mM NaF, 20 mM NEM, 0.25 mM PMSF and 50 U/ml Benzonase (Merck Millipore) and incubated at room temperature for 5 min. Soluble fractions were prepared by rotation at 4°C for 1 hour after adjusting the NaCl concentrations to 300 mM, followed by centrifugation at 20,000 x g for 10 min at 4°C. The protein concentrations of cell extracts were determined with Coomassie Protein Assay Reagent (Thermo Scientific) with bovine serum albumin (BSA) standard (Takara). The antibodies used in this research are described Table S2.

### Immunofluorescence staining

For subcellular localization analysis of USP42, cells were fixed with 4% paraformaldehyde (PFA) for 15 min at room temperature and then permeabilized by incubation with 0.2% Triton X-100 in PBS for 5 min at room temperature. To examine 53BP1 foci formation, cells that were irradiated with 2 Gy of IR and then incubated for 15 min were pre-extracted prior to fixation with pre-extraction buffer [10 mM Pipes (pH 6.8), 3 mM MgCl_2_, 3 mM EDTA, 0.5% Triton X-100, 0.3 M sucrose and 50 mM NaCl] for 5 min on ice. For the purpose of investigating RAD51 and BRCA1 foci formation, cells that were irradiated with 2 Gy of IR and then incubated for 6 hours were pre-extracted with 0.2% Triton X-100 for 1 or 5 min, respectively and then fixed with 3% PFA and 2% sucrose in PBS for 15 min. Hereafter, the samples were washed twice with 0.1% Tween 20 in PBS after each procedure. After incubating cells with blocking buffer A [5% FBS, 0.1% Triton X-100 in PBS] for 30 min, the cells were sequentially incubated with primary antibodies for 1 hour and with secondary antibodies for 30 min diluted in blocking buffer A for USP42 localization and 53BP1 foci formation analysis. For RAD51 and BRCA1 foci formation analysis, blocking buffer B (2% BSA in PBS) was used instead and incubated for 1 hour prior to the incubation with the antibodies. Following nuclei staining with 1 μg/ml of 4’,6-diamidino-2-phenylindole (DAPI) solution for 10 min, the samples were sealed with VECTASHIELD (Vector), and images were taken with a confocal microscope (Leica, TCS SP5) and analysed with a software (Leica, LAS AF). To analyse 53BP1 foci formation, images were taken by IN Cell Analyzer 2000 (GE Healthcare), and then cells were classified into the S, G2 and G1 phases based on the signal intensity of anti-CENPF antibody staining with software (IN Cell Investigator, GE Healthcare).

### Cell cycle profile analysis

The cell cycle profile was analysed with BrdU incorporation as previously described (29).

### Quantitative DNA-end resection assay

The efficiency of DNA-end resection was measured in a quantitative manner, as previously described (29). Briefly, cells were labelled with BrdU for 24 hours prior to 1 μM CPT treatment for 1 hour. The cells were processed for staining with anti-BrdU and anti-Cyclin A antibodies under non-denaturing conditions, followed by incubation with appropriate secondary antibodies, and then analysed with LSRFortessa (BD Biosciences). The signal intensity of the anti-BrdU antibody was obtained from a subpopulation of the cells that were positive for anti-Cyclin A antibody staining by FlowJo software (TreeStar).

### Establishment of the USP42 KO cell line and its complemented cell lines

The gRNA sequence was cloned between the BamHI site and the BsmBI site of the pCas-Guide vector (OriGene). Sequences of the homology arms flanking the PAM site were cloned into pDonor-D09 (GeneCopoeia). U2OS cells co-transfected with these plasmids were cultured for 14 days with puromycin and screened for loss of USP42 expression by immunoblotting. To obtain complemented cell lines, the USP42 KO cell line was transfected with the plasmids encoding GFP or GFP-USP42 (FL, ΔC or Δ946-1196) and cultured for 14 days with 1 mg/ml Geneticin.

### Clonogenic cell survival assay

Clonogenic viability was examined using a colony formation assay. Briefly, forty-eight hours after the initial transfection with siRNAs, cells were seeded in 6-well plates and treated with acute IR on the following day. For the assay with the USP42 KO cell line, cells were seeded in 6 well plates one day before irradiation. Colonies were stained with crystal violet solution [2% crystal violet (Sigma-Aldrich) in 10% ethanol] 10-13 days after IR treatment.

### Immunoprecipitation

Soluble fractions of cell extracts were immunoprecipitated with an anti-GFP antibody coupled to magnetic beads (GFP-Trap_MA, ChromoTek) or an anti-HA antibody coupled to agarose beads (EZview Red Anti-HA Affinity Gel, Sigma-Aldrich) by rotation overnight at 4°C. The beads were washed six times with the buffer used for cell extract preparation and bound proteins were eluted by boiling at 95°C for 10 min with 1 x Laemmli SDS buffer [62.5 mM Tris-HCl (pH 6.8), 2% sodium dodecyl sulfate, 10% glycerol and 0.02% bromophenol blue, 6.25% β-mercaptoethanol and 300 mM NaCl].

### Mass spectrometry analysis

Cell extracts were prepared with the Benzonase-based method (see above) from U2OS cells transiently transfected with pEGFP-C1-USP42 or pEGFP-C1 as a negative control. Immunoprecipitates generated with an anti-GFP antibody were washed six times with IP lysis buffer containing 300 mM NaCl, 1 x PI, 10 mM NaF, 20 mM NEM and 0.25 mM PMSF and then collected by boiling at 95°C for 10 min with 1 x Laemmli SDS buffer. Immunoprecipitated samples were digested in-solution and analysed using nanoliquid chromatography tandem mass spectrometry (nano–LC-MS/MS) provided by Filgen.

### Slot blot analysis for R-loop quantification

Slot blot analysis was basically performed as previously described (54). Briefly, genomic DNA (1 μg) was transferred to a nylon membrane using slot blot apparatus in duplicate for R-loop detection with an S9.6 antibody and DNA staining with ethidium bromide. The signal intensity obtained with an S9.6 antibody was normalized by the signal intensity with ethidium bromide.

### Statistical analysis

All statistical analyses were performed using a standard two-sided Student’s t-test.

## Acknowledgements

We thank Stephen P. Jackson (Cambridge, UK) for helping us to set the research group up. This work was funded by Grant-in-Aid for Research Activity start-up 15H06738 (R.N.), Grant-in Aid for Young Scientists (A) 16H05888 (R.N.), the Daiichi Sankyo Foundation of Life Science (R.N.), the Mochida Memorial Foundation for Medical and Pharmaceutical Research (R.N.), the Takeda Science Foundation (R.N.) and the Uehara Memorial Foundation (R.N.). Research in the K.I. laboratory is supported by Grant-in Aid for Scientific Research (C) 18K11648 (R.S.) and Grant-in Aid for Scientific Research (B) 18H03375 (R.S. and K.I.). Research in the M.T. laboratory is funded by Grant-in Aid for Young Scientists (B) 17K12822 (Y.K.), Grant-in Aid for Scientific Research (C) 18K11642 (M.I.) and Grant-in Aid for Scientific Research (A) 15H01738 (M.I. and M.T.). The Radiation Biology Center is a joint usage research center supported by the MEXT of Japan. A part of the authors’ work has been performed in collaborative research projects in the Joint Usage Research Center.

## Author contributions

Conceptualization, M.M. and R.N.; Formal Analysis, M.M. and R.N.; Investigation, M.M., R.S., M.A., Y.K., S.K., W.T. and R.N.; Resources, Y.K., M.I., K.I. and M.T.; Writing-Original Draft, R.N.; Writing-Review & Editing, M.M., R.S., M.A., Y.K., M.I., K.I., M.T. and R.N.; Funding Acquisition, R.S., Y.K., M.I., K.I., M.T. and R.N.

## Supplemental information legends

**Fig. S1. Nuclear speckle factor screening for HR regulation indicated a cross-talk between HR and transcription. Related to Fig. 1**

(A) U2OS cells were subjected to immunofluorescence staining with anti-USP42 and anti-SC35 antibodies. Nuclei were stained with DAPI. Scale bar: 10 μm.

(B) U2OS cells transfected with the indicated siRNAs were subjected to immunoblotting analysis with the indicated antibodies.

**Fig. S2. USP42 promotes HR by recruiting BRCA1 to DSB sites. Related to Fig. 2**

(A) RAD51 foci formation efficiency was examined with the cells transfected with the indicated siRNAs. The Representative images are shown. Scale bar: 10 μm.

(B) A DR-GFP assay was performed with the indicated siRNAs (mean ± SEM, n=3).

(C) U2OS cells were transfected with the indicated siRNAs and then cell cycle profiles were analysed (mean ± SEM, n=3).

(D) U2OS cells transfected with the indicated siRNAs were subjected to immunoblotting analysis with the indicated antibodies.

(E) U2OS and USP42 KO cells were treated with CPT (1 μM, 1 hour) and analysed by immunoblotting with the indicated antibodies.

(F) U2OS and USP42 KO cells were investigated for the expression of HR factors by immunoblotting with the indicated antibodies.

(G) BRCA1 foci formation efficiency was examined with the cells transfected with the indicated siRNAs. Representative images are shown. Scale bar: 10 μm.

***: p< 0.005.

**Fig. S3. Nuclear speckle localization of USP42 is required for efficient homologous recombination. Related to Fig. 3**

(A) Schematic representation of truncated USP42 proteins. The numbers represent amino acid residues. R: arginine-rich region.

(B) Subcellular localization of GFP-fused truncated USP42 proteins. Cells transfected with the plasmids coding the indicated GFP-USP42 were subjected to fluorescence microscopy analysis. Scale bar: 10 μm.

(C) Amino acid sequence of USP42 (946-1196 a. a.) in which RS repeats are indicated with red characters. The numbers represent amino acid residues.

(D) The probability of disorder of USP42 predicted by IUPred2A was plotted against amino acid residues. A schematic representation of USP42 is also shown underneath the graph. The USP domain (green) and nuclear speckle localization region (red) are indicated.

**Fig. S4. USP42 is epistatic with DHX9 in the cellular survival after DSB induction and promotes resolution of DSB-induced R-loop. Related to Fig. 4**

(A) A DR-GFP assay was performed with the indicated siRNAs (mean ± SEM, n=3).

(B) U2OS cells were transfected with the indicated siRNAs, and then cell cycle profiles were analysed (mean ± SEM, n=3).

(C) The indicated cells were analysed for R-loop with an S9.6 antibody. Signal intensity was normalized to U2OS cells, and then plotted as a box and whiskers plot (median, 5 to 95 percentile, n=4). *: p<0.05.

**Table S1. The oligonucleotides used in this research.**

**Table S2. The antibodies used in this research.**

