## Supplemental Information for "A novel nuclear speckle factor, USP42, promotes homologous recombination repair by resolving DNA double-strand break induced R-loop"

**This PDF file includes**

Figs. S1 to S4

Table S1 and S2

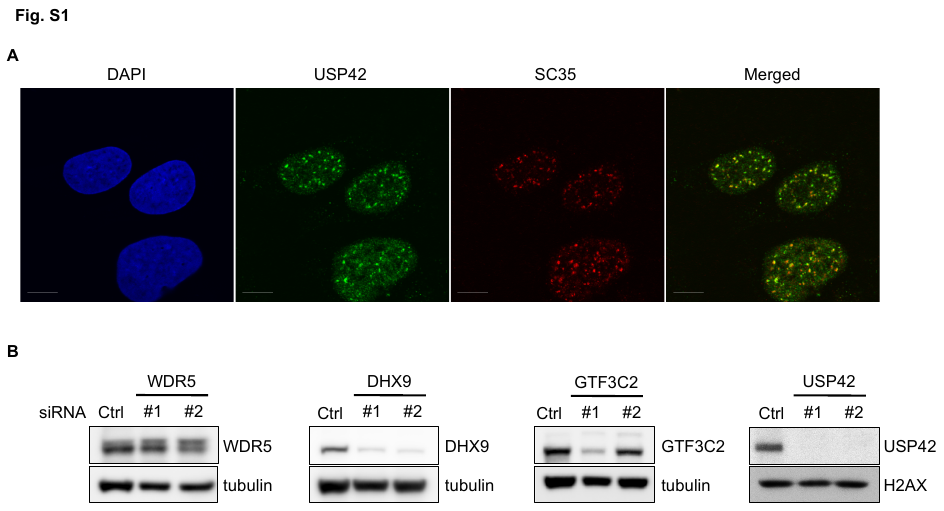

**Fig. S1. Nuclear speckle factor screening for HR regulation indicated a cross-talk between HR and transcription. Related to Fig. 1**

(A) U2OS cells were subjected to immunofluorescence staining with anti-USP42 and anti-SC35 antibodies. Nuclei were stained with DAPI. Scale bar: 10 μm.

(B) U2OS cells transfected with the indicated siRNAs were subjected to immunoblotting analysis with the indicated antibodies.

**Fig. S2. USP42 promotes HR by recruiting BRCA1 to DSB sites. Related to Fig. 2**

(A) RAD51 foci formation efficiency was examined with the cells transfected with the indicated siRNAs. The Representative images are shown. Scale bar: 10 μm.

(B) A DR-GFP assay was performed with the indicated siRNAs (mean ± SEM, n=3).

(C) U2OS cells were transfected with the indicated siRNAs and then cell cycle profiles were analysed (mean ± SEM, n=3).

(D) U2OS cells transfected with the indicated siRNAs were subjected to immunoblotting analysis with the indicated antibodies.

(E) U2OS and USP42 KO cells were treated with CPT (1 μM, 1 hour) and analysed by immunoblotting with the indicated antibodies.

(F) U2OS and USP42 KO cells were investigated for the expression of HR factors by immunoblotting with the indicated antibodies.

(G) BRCA1 foci formation efficiency was examined with the cells transfected with the indicated siRNAs. Representative images are shown. Scale bar: 10 μm.

***: p< 0.005.

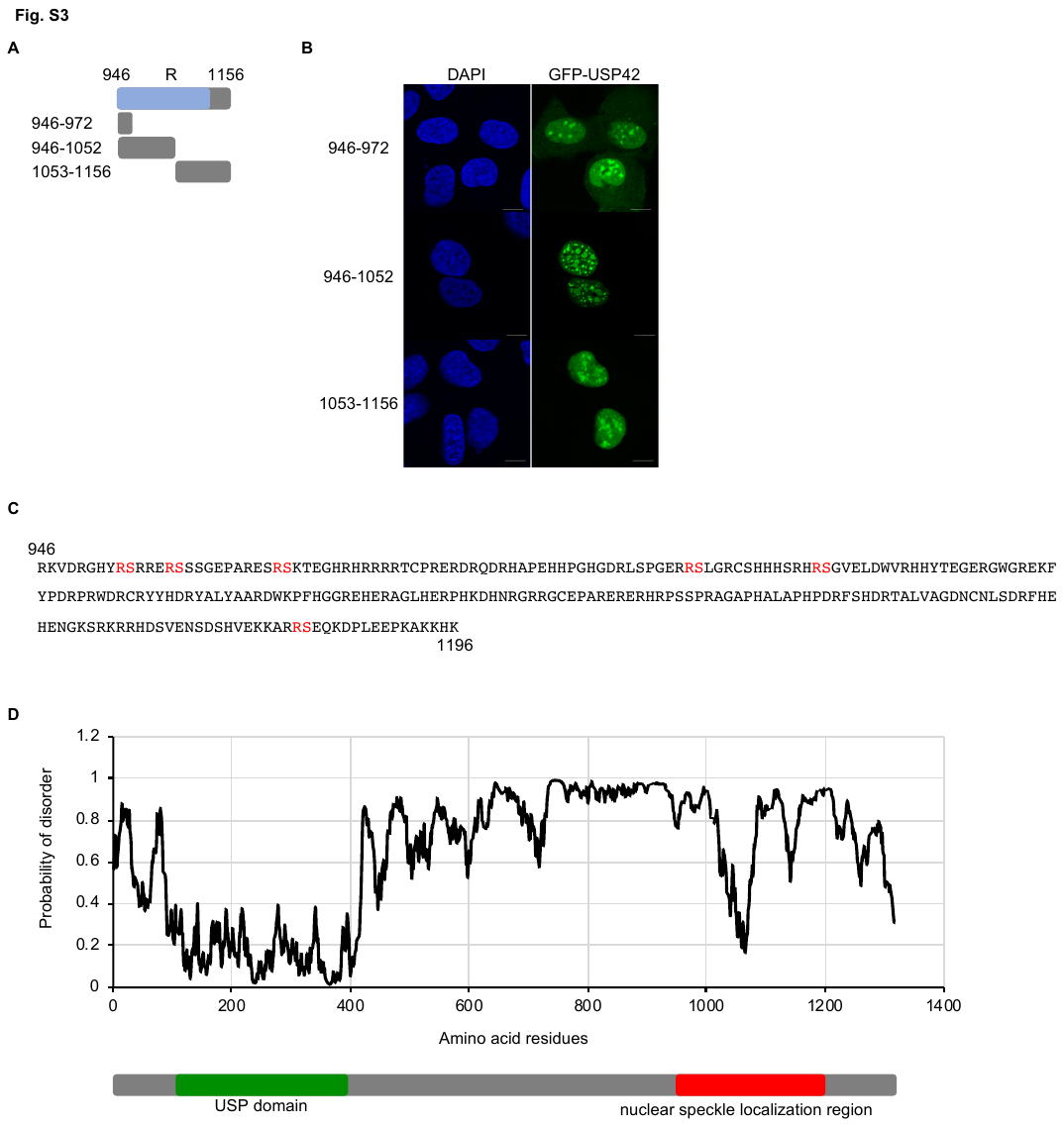

**Fig. S3. Nuclear speckle localization of USP42 is required for efficient homologous recombination. Related to Fig. 3**

(A) Schematic representation of truncated USP42 proteins. The numbers represent amino acid residues. R: arginine-rich region.

(B) Subcellular localization of GFP-fused truncated USP42 proteins. Cells transfected with the plasmids coding the indicated GFP-USP42 were subjected to fluorescence microscopy analysis. Scale bar: 10 μm.

(C) Amino acid sequence of USP42 (946-1196 a. a.) in which RS repeats are indicated with red characters. The numbers represent amino acid residues.

(D) The probability of disorder of USP42 predicted by IUPred2A was plotted against amino acid residues. A schematic representation of USP42 is also shown underneath the graph. The USP domain (green) and nuclear speckle localization region (red) are indicated.

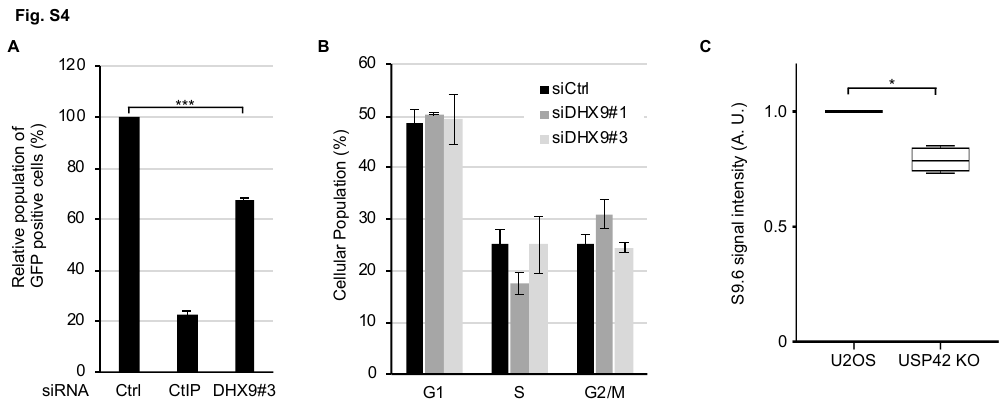

**Fig. S4. USP42 is epistatic with DHX9 in the cellular survival after DSB induction and promotes resolution of DSB-induced R-loop. Related to Fig. 4**

(A) A DR-GFP assay was performed with the indicated siRNAs (mean ± SEM, n=3).

(B) U2OS cells were transfected with the indicated siRNAs, and then cell cycle profiles were analysed (mean ± SEM, n=3).

(C) The indicated cells were analysed for R-loop with an S9.6 antibody. Signal intensity was normalized to U2OS cells, and then plotted as a box and whiskers plot (median, 5 to 95 percentile, n=4). *: p<0.05.

**Table S1. The oligonucleotides used in this research.**

| Oligonucleotides | SOURCE | IDENTIFIER |
| --- | --- | --- |
| siRNA targeting sequence: DHX9#1: CUGCUAUCAUCAGCCAGUU | This paper | N/A |
| siRNA targeting sequence: DHX9#2: CAGGAUAUGUGCAGGUGUU | This paper | N/A |
| siRNA targeting sequence: DHX9#3: GUAAAUGAACGUAUGCUGA | This paper | N/A |
| siRNA targeting sequence: USP42#1: GUUAAUAGGUCCUCAGUGA | This paper | N/A |
| siRNA targeting sequence: USP42#2: CCCUCUUCUACCAUUACCA | This paper | N/A |
| siRNA targeting sequence: USP42#3: GCGUGGUGCUUCUUUAGUA | This paper | N/A |
| siRNA targeting sequence: WDR5#1: GUCCUUGUGAAGCUCGUCU | This paper | N/A |
| siRNA targeting sequence: WDR5#2: GUCGUGAUCUCAACAGCUU | This paper | N/A |
| siRNA targeting sequence: GTF3C2#1: CUGUCAACCAUCACUACUU | This paper | N/A |
| siRNA targeting sequence: GTF3C2#2: GACCAAGGAAUACCACAGA | This paper | N/A |
| siRNA targeting sequence: CtIP: GCUAAAACAGGAACGAAUC | Sartori et al., 2007 | N/A |
| siRNA targeting sequence: Ctrl: AACGUACGCGGAAUACUUCGA | Nishi et al., 2014 | N/A |
| gRNA sequence for knocking out USP42: ATTGGTTTAATAACGTCCCC | This paper | N/A |
| Primer for USP42 left homology arm cloning forward: GGGAATTCCTCATCCCTGCTGTGGGAGACTTTC | This paper | N/A |
| Primer for USP42 left homology arm cloning reverse: GGGGTACCACTGAGTGCCTGGGTAATATGTGCTT | This paper | N/A |
| Primer for USP42 right homology arm cloning forward: GGTCTAGAAATGTTTGTCATCAATGAGATGCGGC | This paper | N/A |
| Primer for USP42 right homology arm cloning reverse: GGGTCGACGGCAGTGTGTCCAGAGTAGCCTAAT | This paper | N/A |
| siRNA targeting sequence: DHX9#1: CUGCUAUCAUCAGCCAGUU | This paper | N/A |
| siRNA targeting sequence: DHX9#2: CAGGAUAUGUGCAGGUGUU | This paper | N/A |
| siRNA targeting sequence: DHX9#3: GUAAAUGAACGUAUGCUGA | This paper | N/A |
| siRNA targeting sequence: USP42#1: GUUAAUAGGUCCUCAGUGA | This paper | N/A |
| siRNA targeting sequence: USP42#2: CCCUCUUCUACCAUUACCA | This paper | N/A |
| siRNA targeting sequence: USP42#3: GCGUGGUGCUUCUUUAGUA | This paper | N/A |
| siRNA targeting sequence: WDR5#1: GUCCUUGUGAAGCUCGUCU | This paper | N/A |
| siRNA targeting sequence: WDR5#2: GUCGUGAUCUCAACAGCUU | This paper | N/A |
| siRNA targeting sequence: GTF3C2#1: CUGUCAACCAUCACUACUU | This paper | N/A |

**Table S2. The antibodies used in this research.**

| Antibodies | SOURCE | IDENTIFIER |
| --- | --- | --- |
| Anti-WDR5 antibody | Abcam | Cat#ab22512 |
| Anti-RNA Helicase A antibody | Abcam | Cat#ab54593 |
| GTF3C2 Antibody | Proteintech | Cat#27494-1-AP |
| Anti-USP42 antibody produced in rabbit | ATLAS | Cat#HPA006752 |
| Anti-USP42 antibody | Abcam | Cat#ab121254 |
| Monoclonal Anti-splicing Factor SC35 antibody produced in mouse | Sigma-Aldrich | Cat#S4045 |
| Anti-SC35 antibody-Nuclear Speckle Marker | Abcam | Cat#ab11826 |
| Anti-RPA32/RPA2 (phosphor S4+S8) antibody | Abcam | Cat#ab87277 |
| Anti-phospho-Histone H2A.X (Ser139) antibody, clone JBW301 | Merck Millipore | Cat#05-636 |
| Monoclonal Anti-α-Tubulin antibody produced in mouse | Sigma-Aldrich | Cat#T9026 |
| Anti-BrdU Antibody | GE Healthcare | Cat#RPN202 |
| Anti-RAD51 Antibody | Bio Academica | Cat#70-001 |
| BRCA1 antibody (D-9) | Santa Cruz Biotechnology | Cat#sc-6954 |
| Anti-53BP1 Antibody, clone BP13 | Merck Millipore | Cat#MAB3802 |
| Anti-RPA32/RPA2 antibody [9H8] | Abcam | Cat#ab2175 |
| Anti-Histone H2A.X antibody-ChIP Grade | Abcam | Cat#ab11175 |
| Cyclin A (H-432) Antibody | Santa Cruz Biotechnology | Cat#sc-751 |
| Mre11 antibody [12D7] | Gene Tex | Cat#GTX70212 |
| Rad50 antibody [13B3] | Gene Tex | Cat#GTX70228 |
| Anti-p95/NBS1 antibody | Abcam | Cat#ab23996 |
| CtIP antibody (mAb) | Active Motif | Cat#61141 |
| Anti-CENPF antibody | Abcam | Cat#ab5 |
| Anti-GFP | Roche | Cat#11814460001 |
| Ubiquitin (P4D1) Mouse mAb | Cell Signaling TECHNOLOGY | Cat#3936S |
| Anti-HA (12CA5) | Roche | Cat#11583816001 |
| Anti-DNA-RNA Hybrid Antibody, clone S9.6 | Merck Millipore | Cat#MABE1095 |
